# *DateBack*, an evolving open-access repository of *Phoenix* archaeobotanical data supporting new perspectives on the history of date palm cultivation

**DOI:** 10.1101/2025.02.21.639487

**Authors:** Margot Besseiche, Elora Chambraud, Vladimir Dabrowski, Elisa Brandstätt, François Sabot, Charlène Bouchaud, Muriel Gros-Balthazard

## Abstract

The date palm (*Phoenix dactylifera* L.) has been a cornerstone of oasis agrosystems in West Asia and North Africa for millennia, yet the timeline and processes of its domestication and spread remain poorly documented. Archaeobotanical remains provide critical insights into its cultivation history, but no comprehensive review or dedicated platform currently exists to synthesize and analyze these data.

To address this gap, we conducted an extensive literature review and developed *DateBack* (https://cloudapps.france-bioinformatique.fr/dateback), a digital open-access repository of archaeobotanical remains of *Phoenix* (L.) spp. In its first version, *DateBack* records macroremains (e.g., seeds, fruits, stems, petioles) from prehistoric to Late Antiquity contexts in Southwest and South Asia.

We assembled 154 entries from 110 archaeological sites across 123 references, along with a separate table of 74 radiocarbon-dated entries from 20 sites, refining chronological assessments. Most entries comprise charred seeds (58.4%), with a smaller proportion of charred vegetative parts or desiccated remains. Our findings highlight regional disparities in the distribution of remains, with concentration in the Levant and the Gulf region, while Saudi Arabia and southern Iran are underrepresented. There are also disparities in data reliability, particularly in dating resolution, which complicate the reconstruction of date cultivation history. Our evidence shows that the earliest securely dated macro-botanical remains, indicating date consumption, appear in the Gulf region around 5000 BCE, while cultivation emerges later, first in southern Mesopotamia and the northern Gulf in the 4^th^ millennium BCE, then in eastern Arabia and the Indus Valley in the 3^rd^ millennium BCE. The Levant presents challenges, with numerous presumed early finds but significant uncertainties, with secure evidence of cultivation only appearing by the late 2^nd^ millennium BCE, as in northwestern Arabia, while in the southern Arabian Peninsula, it is not attested until the 1^st^ millennium BCE.

By compiling and standardizing archaeobotanical data, *DateBack* facilitates advanced spatiotemporal analyses of date palm history and supports additional archaeobotany-based research including ancient DNA studies. Moreover, the platform is dynamic, scalable, and collaborative, enhancing data integration and refinement, with future expansions planned to include additional periods (Islamic era and beyond), geographic regions (North Africa), and new types of evidence, such as plant microremains and iconography.

## Introduction

The date palm *(Phoenix dactylifera* L.) is a keystone species in oasis agrosystems of North Africa and West Asia, where it plays a crucial role in agricultural activities, human livelihoods, and cultural traditions. Cultivated in more than 30 countries today, the date palm yields over 10 million tons of sugar-rich dates annually, a figure that continues to grow (FAOSTAT, 2023). Beyond its role as a dietary staple, all parts of the plant have long been used for diverse purposes: leaves and fibers for making baskets and ropes, as well as the stem and petiole for building (as beam, roof) and fuel use, especially in wood-scarce environments (e.g., Mouton et al., 2012; Tengberg, 2012). Its canopy and surroundings further support the cultivation of other crops, such as cereals, pulses, herbs, legumes, and fruit trees, in arid environments (Tengberg, 2012). The cultural and traditional significance of the date palm is recognized internationally, as reflected in the 2022 UNESCO inscription of its associated knowledge, skills, and practices on the Representative List of the Intangible Cultural Heritage of Humanity.

While the date palm’s significance in agriculture and culture is well-established, critical aspects of its early history remain unclear, including its origins, routes and timing of spread, and its role in shaping oasis agriculture (reviewed in Gros-Balthazard & Flowers, 2021). *Phoenix dactylifera* L. (1753) belongs to the genus *Phoenix* L., which comprises 12 other non-cultivated species (Barrow, 1998; Gros-Balthazard et al., 2021; Cid-Vian et al., 2025). Although genetic studies confirm that domestication stemmed from *P. dactylifera* itself (Pintaud et al., 2010; Gros-Balthazard et al., 2017), the prehistoric distribution of its ancestral populations remains poorly understood, hindering efforts to fully reconstruct its early interactions with human societies (Gros-Balthazard & Flowers, 2021).

Archaeological evidence points to date consumption as early as ∼5000 BCE in the Gulf region, based on date seeds recovered from ancient deposits (Beech & Shepherd, 2001; Parker, 2010). By the late 4^th^ millennium BCE, archaeobotanical, written and iconographic evidence indicate widespread date palm cultivation in Mesopotamia, leading some researchers to propose it as a domestication center (Tengberg, 2012; Zohary & Spiegel-Roy, 1975; Landsberger, 1967). Alternatively, others suggest that fruit tree horticulture, including date palm, may have first developed in the Chalcolithic Levant (Zohary et al., 2012; Langgut & Sasi, 2023; Langgut, 2024).In North Africa, date palm cultivation is attested later, with evidence from the 2^nd^ millennium BCE in Egypt (Tengberg & Newton, 2016), early 1^st^ mill. BCE in the central Sahara (Van der Veen, 1992; Kaczmarek et al., 2024), and only as late as medieval times in Morocco (Ros et al., 2024).

While this pattern in the archaeological data suggests a westward diffusion of date palm cultivation from West Asia across its historical range, spanning Morocco to Pakistan (Tengberg, 2012), genetic data challenge this straightforward narrative. Cultivated date palms are divided into two primary genepools — North African and West Asian (Zehdi-Azouzi et al., 2015; Hazzouri, Flowers et al., 2015) — which some researchers interpret as evidence of two distinct domestication centers (Zehdi-Azouzi et al., 2015). North African date palms exhibit higher genetic diversity than would be expected from a simple diffusion model (Hazzouri, Flowers et al., 2015; Gros-Balthazard et al., 2017). This unexpected diversity may partly be explained by inter-specific hybridization with *Phoenix theophrasti* Greuter, 1967, a wild relative currently restricted to Crete and Turkey, though the timing and mechanisms of such hybridization events remain unclear (Flowers et al., 2019; Gros-Balthazard, Battesti al., 2020; Gros-Balthazard et al., 2021; Pérez-Escobar, Bellot et al., 2021). Together, these findings highlight a complex history of spread and diversification, requiring further investigation to untangle the interactions between migration, hybridization, and local adaptation.

Archaeobotanical disciplines bring major insights to understand the origins, domestication, diffusion and diversification of crops. Plant macro-(wood, seeds, fruits, and other non-woody remains) and microremains (pollen, phytoliths, starch) recovered from archaeological sites provide insights into several aspects of plants-human interactions of past societies such as plant domestications, subsistence strategies, agropastoral practices, crop-processing, or trade and craft activities (Cappers & Neef, 2012; Marston & Srour, 2014; Van der Veen, 2018). Macroremains are particularly valuable for reconstructing crop domestication and diffusion patterns. They are widely available on archaeological sites, their diverse modes of conservation (e.g., as charred, mineralized, waterlogged, desiccated items) increases their likelihood of being preserved, and they allow more accurate crop identification compared to other botanical proxies, such as phytoliths. For example, identifying phytoliths at a low taxonomic level is challenging, as the morphological similarities within the Arecaceae family make it difficult to differentiate *Phoenix* from other taxa (e.g., Bretzke et al., 2013). Some anatomical parts (e.g., seeds or fruits) exhibit traits associated with domestication syndrome (Cappers & Neef, 2021; Fuller, 2018). In particular, seed morphometrics can be used to differentiate between wild and cultivated accessions and thus trace the emergence of domestication and cultivation (e.g., in grape, Terral et al., 2010; date palm, Gros-Balthazard et al., 2016). However, depending on the context, the presence of macroremains (e.g., seeds) can suggest consumption rather than cultivation (Van der Veen, 2011). While fruits and seeds can travel with humans and may not necessarily be of local origin, the presence of vegetative parts, such as leaves or stems, suggests a higher likelihood of local cultivation. Additionally, desiccated macroremains recovered from arid environments are common (Celant et al., 2015; Van der Veen, 2007; Malleson & Srour, 2025) and provide a suitable material for ancient DNA analysis to better understand patterns of genetic diversity and evolutionary change (e.g., in date palm, Pérez-Escobar, Bellot et al., 2021). However, this state of preservation may raise issues when it comes to precise dating since there is a possibility of modern contamination (e.g., Kaczmarek et al., 2024).

Despite the potential of archaeobotanical data to reconstruct crop history, several challenges persist. A significant amount is scattered across grey literature, excavation reports, and peer-reviewed publications, with inconsistencies in reporting, lack of standardization (e.g., Bates & Jiménez-Arteaga, 2024), and barriers posed by diverse formats and languages. Efforts to compile archaeobotanical information, such as ArchbotLit (Kirleis et al., 2021), provide valuable lists of publications mentioning specific taxa, including the date palm. However, these tools do not integrate datasets or visualization capabilities, limiting their usefulness for broader analyses. While other platforms aim to create structured databases for archaeobotanical data, such as ADEMNES (https://www.ademnes.de/), ArboDat+ (http://arbodat.info/), I2AF (https://i2af.mnhn.fr/), they often face similar issues, including incomplete coverage or a lack of analytical tools. Yet, meta-analyses of archaeobotanical remains have proven invaluable for uncovering the history and processes of crop domestication and dispersal (e.g., Fuller, 2018; Silva et al., 2015; Terral et al., 2010), highlighting the significant opportunities that such integrative resources can unlock.

Among crops, these challenges are especially marked for the date palm, where existing reviews, while valuable, remain narrow in scope. Focused on specific geographic areas (e.g., the Jordan Valley, Langgut, 2024; or the Persian Gulf, Tengberg, 2012) or particular types of remains (e.g., seeds, Rivera et al., 2014), these studies often lack the standardization and comprehensive datasets required for broader syntheses. As a result, gaining insights from archaeobotanical data into the history, domestication, and diffusion of the date palm continues to be a major challenge. This difficulty is further compounded by the need to reassess published data in light of recent advances—such as the identification of domestication syndrome and improvements in taxonomic resolution (e.g., Thomas, 2013; Gros-Balthazard et al., 2016; 2017)—which further underscore the importance of developing integrated and updatable tools for data synthesis.

In this study, we aim to address these gaps by developing a centralized and evolving web platform that compiles archaeobotanical evidence for the date palm and its wild relatives (*Phoenix* spp.). The first version of *DateBack*, presented here, focuses on macroremains from pre-Islamic periods, spanning prehistory to Late Antiquity, in West and South Asia, creating a dedicated repository based on an extensive literature review. The platform consolidates data with a strong emphasis on methodological rigor, verification of the original sources, standardization, contextual details, and integration of radiocarbon information.

By providing tools for visualization and spatial analysis, *DateBack* facilitates archaeobotanical syntheses, allowing researchers to identify patterns and trends in the data. In the present article, we provide a first interpretative synthesis based on the compiled dataset, focusing on the early development of date palm cultivation in different regions of Asia. This preliminary analysis highlights spatial and chronological patterns in the archaeobotanical record, and aims to stimulate further discussion and refinement as the platform evolves. Furthermore, *DateBack* lays the foundation for future interdisciplinary applications, including ancient DNA studies and seed morphometric analyses, which may provide additional insights into the domestication, cultivation, and diffusion of the date palm. As a dynamic and collaborative resource, it contributes to a broader understanding of early agriculture in arid and semi-arid environments and enhances access to archaeobotanical data for both researchers and the wider public.

## Material and methods

### Scope of the literature review

The review focuses on archaeobotanical remains attributed to the genus *Phoenix*, encompassing all species mentioned in the original sources. This inclusive approach acknowledges the challenges of species-level identification due to morphological and anatomical similarities within the genus (e.g., Thomas, 2013; Gros-Balthazard et al., 2016), as well as the potential contributions of wild relatives to the evolutionary history and diversity of cultivated date palms (e.g., Flowers et al., 2019).

Given the vast number of studies mentioning *Phoenix* remains across its entire distribution range and throughout prehistoric and historical periods, we restricted the scope to work with a manageable subset of data. The first iteration of the *DateBack* platform was thus designed to focus on the emergence and early diffusion of date palm cultivation, which was achieved by applying restrictions based on material type, geographical focus, and chronology:

- The review focused on macroremains, including seeds and fruits, inflorescence (flower, perianth, and pedicel), and vegetative parts (stem, petiole, leaf/leaflet, fibre, root). Microremains, including phytoliths, were excluded from the review due to the difficulties in confidently attributing them to *Phoenix*, as they are often indistinguishable from those of other members of the Arecaceae family (e.g., Bretzke et al., 2013). Alongside with pollen, they are usually sampled on off-site, non-anthropogenic locations (e.g., palaeolakes, stalagmites) for the purpose of palaeoenvironmental reconstructions, which fall outside the study (e.g., Parker et al., 2004). Additionally, examples of pollen analysis from archaeological contexts are particularly limited (e.g., Bellini et al., 2011). It is noteworthy, however, that an increase in date palm pollen ratios (or Arecaceae phytoliths) can serve as a reliable indicator of human activity, particularly when associated with archaeological and other archaeobotanical evidence (Langgut, 2024).
- Geographically, the review concentrated on Southwest Asia and part of South Asia, covering the Levantine region, the Arabian Peninsula, and extending eastward across the Indo-Iranian borderlands up to the Indus Valley region. This focus was motivated by two reasons: first, Southwest Asia is widely considered as the region where date palm cultivation emerged (Tengberg, 2012; Langgut & Sasi, 2023), whereas in North Africa, it developed later (Tengberg & Newton, 2016). Second, limiting the scope to Southwest Asia in this first version allowed for a more focused review, given the extensive literature available for North Africa, particularly Egypt.
- Chronologically, the review focused on remains dating up to 630 CE, marking the early emergence of Islam. This cut-off was chosen to manage the large volume of literature and to prioritize the early emergence, diffusion and development of phoeniciculture. By this time, the date palm was cultivated across Southwest Asia and part of South Asia at a broad regional scale (Tengberg, 2012), although new cultivation areas may have continued to emerge.

### Methodology of the literature review

The literature review included a wide range of sources documenting *Phoenix* macroremains, encompassing peer-reviewed journal articles, books, and grey literature such as PhD theses and archaeological excavation reports. For this first version, only references published before November 2024 were considered.

The review began by compiling documents mentioning *Phoenix* remains, using existing literature reviews (e.g., Tengberg, 2012; Flowers et al., 2019: Dataset S1) and other studies referencing such remains (e.g., Weiss, 2015; Langgut & Sasi, 2023). Rather than relying solely on these secondary mentions, we actively sought the original papers documenting the remains. When original papers could not be located, or when they were written in languages not mastered by any of the co-authors, these references were excluded. Particular attention was given to studies documenting early *Phoenix* remains and to key studies from underrepresented regions.

The search was then expanded in two ways. First, we leveraged existing knowledge of excavated sites within the surveyed region, drawing from archaeological databases (e.g., ArchbotLit, Kirleis et al., 2021), excavation reports, insights shared by researchers and archaeobotanists, and bibliographies from articles identified in the initial phase. Second, systematic searches were conducted on Google Scholar, Academia, and the Web of Science using the query “*Phoenix*” to identify documents potentially mentioning archaeobotanical remains. Studies were thoroughly examined and included in the dataset only if they fell within the review’s defined scope.

The objective of this compilation was to be as exhaustive as possible within the defined scope. Particular emphasis was placed on rigorously verifying and critically assessing the occurrences of date palm remains to ensure the reliability of the dataset.

### Data table structure and organization

Based on the review and existing archaeobotanical remains databases, and in line with the objectives of our study, we identified the information to recover (fields), structured the data accordingly, and considered how to standardize this information. The details of this process are described in the results section (“Structure and content of the database”). The data tables are stored as CSV files.

### Web application and visualization tools

The web application and visualization tools were developed using R (v. 4.4.0, R Core Team, 2022) and the shiny package v.1.9.1 (Chang et al., 2012) to facilitate user interaction with the data. These tools were designed to allow users to explore the data table, visualize trends and distributions, and generate interactive maps. The following R packages were used: leaflet v.2.2.2 (Cheng et al., 2015), plotly v.4.10.4 (Sievert, 2020), tidyverse v.2.0.0 (Wickham et al., 2019), viridis v.0.6.5 (Garnier et al., 2023), bslib v.0.8.0 (Sievert et al., 2021), leaflegend v.1.2.1 (Roh, 2024), markdown v.1.13 (Xie et al., 2012). These codes are in open access under GNU GPL3 on a git repository (https://forge.ird.fr/diade/besseiche/dateback).

The web application is hosted by the Institut Français de Bioinformatique (IFB), at https://cloudapps.france-bioinformatique.fr/dateback.

## Results

### Structure and content of the database

The database was built from a systematic review of literature mentioning *Phoenix* archaeobotanical remains, consolidating previously scattered data into a structured format. It was designed to compile and organize key information necessary for both assessing data reliability and enabling visualization and interpretation. It includes details on archaeological sites, chronological contexts, remains data (e.g., taxon, anatomical parts, preservation state), and radiocarbon dating records performed on date palm remains.

The gathered information was organized into two complementary data tables. The structure and detailed descriptions of the main table are provided in Appendix 1. This main table (Appendix 2) serves as the primary repository, capturing detailed information about archaeobotanical records of *Phoenix*. The second table focuses exclusively on radiocarbon-dated date palm remains (Appendix 3). Depending on the level of detail available for each field, some were standardized to finite lists (e.g., “type of archaeological site”), while others were left as free text to capture nuanced or unique information (e.g., “archaeological context”) (Appendix 1).

Briefly, the main data table encompasses the following categories of information:

- Archaeological site details: Name of the site, modern country, type of site (e.g., burial area, settlement), geographic coordinates, and excavation years.
- Chronological context: Description and dating of the context in which the remains were found. Depending on the site, on the number of distinct periods represented, and on the precision of the descriptions in the articles, the context may refer to one stratigraphic unit, locus, area, or several. As a result, multiple “chronological context” sections may exist for the same “archaeological site” section. Information on the nature of the context, including its identification number (if available) or other relevant details, are filled out. The start and end dates represent the earliest and latest possible dates for the context. Broad cultural period and dating quality of the context (whether relative, absolute on material other than date palm, or absolute directly on date palm remain) are also provided when available. For cases of absolute dating on date palm remains, detailed information is provided in the dedicated radiocarbon data table.
- Remains: Type(s) of remains (e.g., seed, stem, petiole), preservation state (e.g., charred, desiccated), taxon as reported in the original paper, and, if necessary, our correction of the taxon with an explanation (see Appendix 1 for details). The number of remains is included as free text to accommodate entries such as “10 seeds,” “>10 seeds,” or “1 seed fragment + unquantified fruits”, reflecting a qualitative approach to quantification, as the complexities and debates surrounding standardization in archaeobotanical studies are beyond the scope of this paper (e.g., Bates & Jiménez-Arteaga, 2024).
- References: The original reference(s) reporting the remain(s) in question, as well as additional sources where other critical information was gathered (e.g., follow-up radiocarbon dating).
- Versioning: Version in which the entry was added and potentially modified, in order to track updates between database versions.

Each entry in the main table represents a unique combination of an archaeological site, contexts with similar start and end dates, and a taxon. To clarify:

- Chronological grouping: Contexts at a given site with similar start and end dates were grouped into a single entry, with identification numbers or other details recorded in the field “chronological context”. For example, at Dalma Island DA11, two contexts (contexts 4 and 15) dated to the same period both yielded date palm remains and were therefore combined into a single entry (Beech & Shepherd, 2001). Conversely, when contexts corresponded to distinct periods, they were recorded as separate entries. For instance, for the site of Qal’at al-Bahrain, four entries were created for four distinct periods: the early, middle, and late Dilmun periods (2200–1750 BCE, 1450–1350 BCE, and 1000–500 BCE, respectively) and the Tylos period (300 BCE–300 CE) (Tengberg & Lombard, 2001). In some cases, overlapping contexts were kept separate when one offered more specific dating within a broader timeframe, as seen at Mleiha (Mouton et al., 2012; Dabrowski, 2019; Tengberg, 1999a; Peña-Chocarro & Barrón Lopez, 1999). Additionally, some contexts were retained as distinct entries despite close date ranges when they corresponded to different archaeological areas. For instance, at Khirbat al-Jariya, two entries have very similar dating (1100 to 900 BCE and 1150 to 900 BCE), which were kept separate since they correspond to a stone structure and a slag mound, respectively (Liss et al., 2020; Stroth et al., 2023; Ben-Yosef et al., 2010).
- Taxon-specific entries: Entries were also categorized based on the taxon reported in the study. When different studies documented distinct taxa at the same site, separate entries were created to reflect these variations. For example, at Jericho, one entry records *Phoenix dactylifera* seeds (Hopf, 1969; Hopf, 1983), while another records charcoal identified as either *Phoenix* or *Hyphaene* (Western, 1971), which was thus classified under Arecaceae.

In addition to the main data table, a complementary table was created to compile radiocarbon data on date palm remains. This so-called ^14^C date palm table includes fields identical to those in the main table to link each radiocarbon analysis to its corresponding site and context. It also contains specific fields such as laboratory codes, anatomical parts dated and their preservation states, raw ^14^C ages (with associated uncertainty), and calibrated age ranges. Versioning information are also provided. Multiple radiocarbon analyses performed on the same site or context are recorded as separate entries to ensure comprehensive documentation.

### Development of the web application and visualization tools

The visualization application consists of a website with two visualization tabs: a *Data Tables* tab which allows the navigation in and the download of the data tables and an *Interactive Map* tab (Figure 1). Both can be interactively explored thanks to various filters. In particular, we provide a filter on the chronological range to explore a given time period. We also allow users to select on the present-day countries they wish to explore, the anatomical parts, their preservation states, the dating qualities of the context and the taxa.

**Figure 1.**
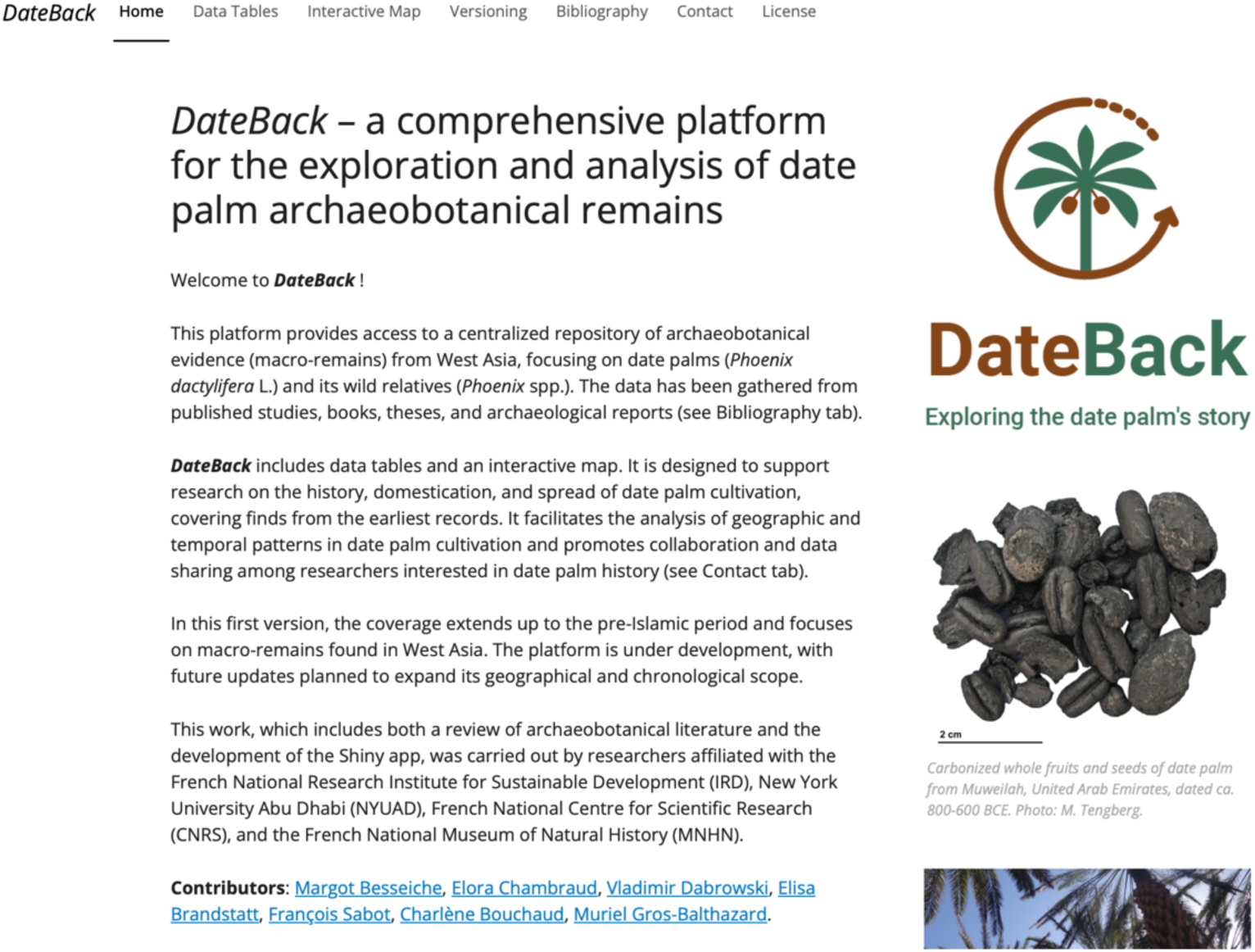
Overview of the *DateBack* platform Homepage. Screenshot of the homepage showcasing the main features of the *DateBack* web platform, including options for data exploration, visualization, and user interaction.

In addition, four informative tabs (*Versioning*, *Bibliography*, Contact and *License*) are accessible. The Contact tab was added to encourage community engagement, thanks to a dedicated email address, for contributions or questions about *DateBack*.

### Results of the literature review

Our literature review encompassed 123 accessible documents that report *Phoenix* spp. remains from prehistoric to pre-Islamic Southwest Asia and South Asia, all of which have been included in the first iteration of the *DateBack* database (v. 1). While we systematically relied on original sources to ensure accuracy, two exceptions were made. First, for the numerous studies authored by Liphschitz in Hebrew, we instead referenced a comprehensive English-language review by the same author, which consolidated detailed information on the sites, contexts, and archaeobotanical findings (Liphschitz, 1996). Second, for two Iranian sites—Tappeh Yaḥyā (Tepe Yahya) and Tepe Gaz Tavila (R37)—we relied on information from Costantini (1985), which constitutes the sole available source of information regarding these sites as the original report by Costantini & Costantini Biasini, announced as “in press” in this article, appears to have never been published.

The main data table includes 154 entries from 110 sites across 15 present-day countries (Appendix 2). The ^14^C date palm datatable contains 74 entries from 20 sites across 7 countries (Appendix 3).

In the original sources, the majority of remains were identified as date palm (*Phoenix* (cf.) *dactylifera*), accounting for 138 entries (89.6%), with only a single report of other species, namely *Phoenix theophrasti*. When original sources lacked precise identification, remains were classified more broadly in the database as *Phoenix* sp. (n=12, 7.8%) or as Arecaceae when identified as either *Phoenix* or another palm genus (n=3, 2.1%). In addition to this compilation of the taxa as reported in the source, we also provided taxonomic corrections in cases of uncertainty. Specifically, for three entries from the Levant, at ‘Atlit (Atlit Yam) and Ohalo II, where both *P. theophrasti* and *P. dactylifera* had been reported, we reclassified them as *Phoenix* sp. to reflect the uncertainty (Galili et al., 1993; Liphschitz & Nadel, 1997; Kislev et al., 2004).

Of the 154 recorded entries, 57 (37.0%) reported multiple anatomical parts of *Phoenix*. The majority of entries comprise seeds (n=113, 73.4%; Figure 2A), a pattern influenced by taphonomic, analytical, and disciplinary biases. Taphonomic factors favor seed preservation due to their durability and resilience, with *Phoenix* seeds in particular demonstrating exceptional resistance to charring (Ivorra et al., 2024). Analytical biases arise because seeds are more easily identifiable at the species or genus level compared to other anatomical parts, owing to their distinct morphological features and the availability of extensive reference collections and seed atlases for taxonomic identification (Nesbitt et al., 2003; Cappers & Neef, 2012). Disciplinary biases further shape the types of remains identified. For example, stems are frequently reported from Levantine sites, reflecting the research emphasis of Liphshitz, whose numerous studies in the region account for half of these entries, illustrating how regional expertise and research specializations influence the types of remains reported.

**Figure 2.**
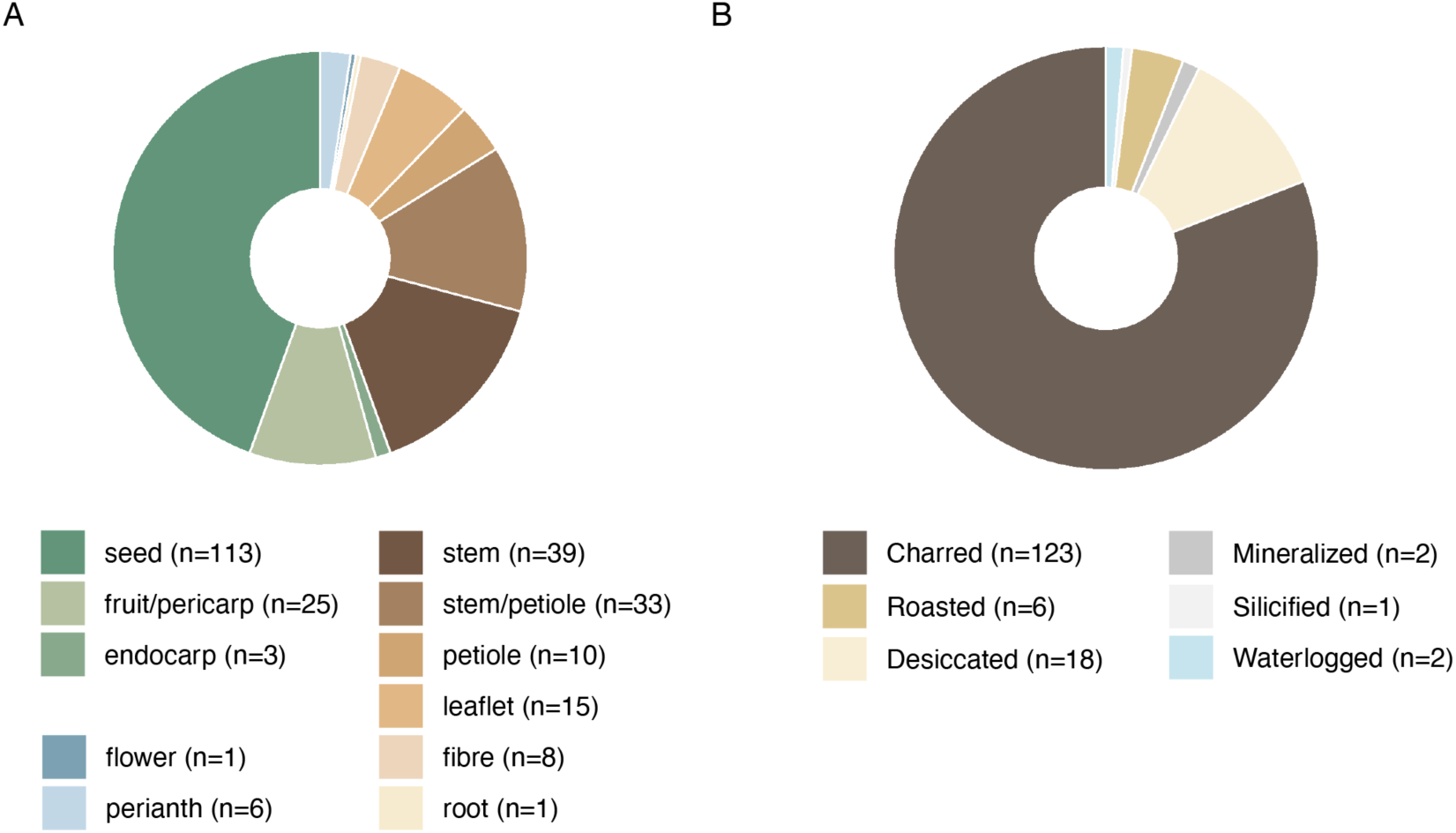
Overview of *Phoenix* archaeobotanical remain from 123 reviewed sources). A. Distribution by type of remains. B. Distribution by preservation state. Since one entry may include several types of remains and multiple preservation states, the total count exceeds the number of entries in the dataset.

Most entries comprise charred remains (n=123, 79.9%, Figure 2B), which aligns with expectations given the prevalence of burning practices and the preservation conditions typical of many archaeological contexts (Van der Veen, 2007). However, the discovery of some desiccated remains (n=18, 11.7%) is particularly noteworthy, as their preservation provides unique opportunities for ancient DNA studies (e.g., Pérez-Escobar, Bellot et al., 2021). Only two entries report waterlogged remains, both from the submerged site of ‘Atlit in the Mediterranean (Galili et al., 1993; Kislev et al., 2004). As expected, such remains are rare in arid West Asia, unlike in Europe, where waterlogged remains, such as grape, are more commonly found (Colledge & Connolly, 2014; Bouby et al., 2023).

The earliest recorded *Phoenix* remains are from Ohalo II in the Levant (21000 BCE to 17000 BCE, Liphshitz & Nadel, 1997), followed by a long gap until approximately 10000 years ago, with remains only becoming more frequent after 5000 BCE, steadily increasing through the 4^th^ millennium BCE to the pre-Islamic period (up to 630 CE) (Figures 3-4). Half (n=76) of the entries come from contexts with radiocarbon dating, either from layers that are radiocarbon-dated but not necessarily the date palm remains themselves (n=49) or from directly radiocarbon-dated date palm remains (n=27), while the rest are either relatively dated or lack chronological information.

**Figure 3.**
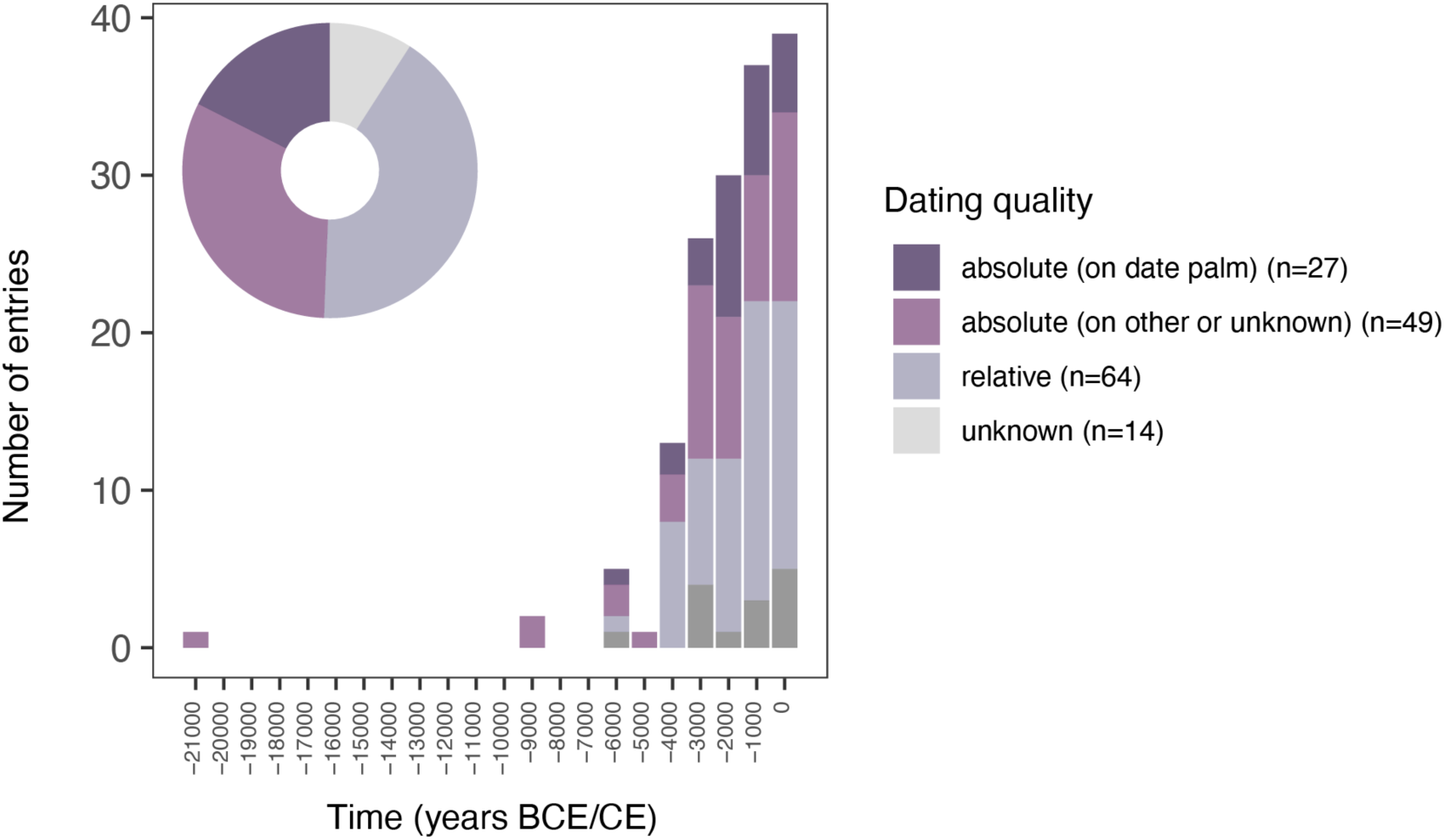
Chronological distribution of *Phoenix* remains from the 123 reviewed sources. Number of entries in the main data table categorized by millennium. The pie chart on the top left represents the total proportions of dating qualities.

**Figure 4.**
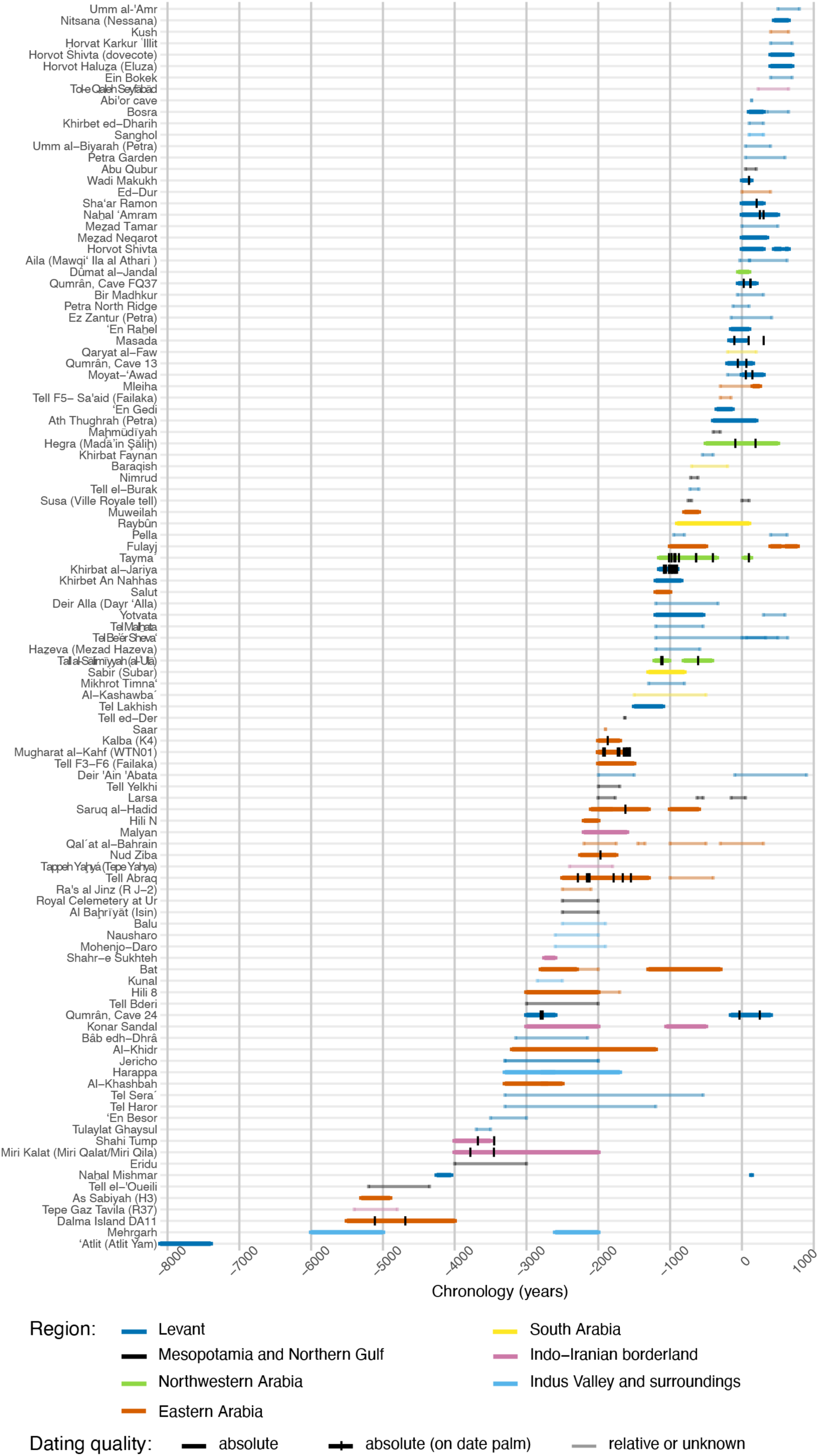
Chronological distribution of the 110 archaeological sites with *Phoenix* macroremains reported in the 123 reviewed sources. The name of the sites are color-coded according to their geographic region. Archaeological contexts with absolute dating are represented by wide, opaque bars, while those with relative or unknown dating quality are shown with thinner, more transparent bars. For contexts with absolute dating directly derived from date palm remains, a vertical line indicates the median calibrated date. These dates were obtained from uncalibrated radiocarbon dates from date palm remains (Appendix 3) and calibrated using the IntCal20 calibration curve (Reimer et al., 2020), implemented in R via the *rcarbon* package (Crema & Bevan, 2021). The vertical line represents 1950 minus the median calibrated BP value, yielding the calibrated BCE date.

Geographically, date palm remains are concentrated in certain regions, while other areas are notably lacking (Figures 4-5). Although this partly reflects actual distribution patterns, the disparity may also result from excavation biases. For example, the Levant has a long history of excavations that is reflected in the results with 68 entries (44.2%). Conversely, although Saudi Arabia covers a third of the surface of the studied region, solely nine entries (5.8%) are from this country, with solely five archaeological sites with published recorded archaeobotanical remains; this will change as ongoing archaeological projects in Saudi Arabia continue to expand (Dabrowski, Bouchaud et al., 2024). Additionally, biases in the retrieval of archaeobotanical remains can arise from early excavation priorities, such as in Iraq, where the focus was predominantly on monumental architecture and artifacts, leaving plant remains largely uncollected or unstudied. This trend was common in the early 20^th^ century, when archaeobotany was not yet a well-established field (Zohary et al., 2012), resulting in significant gaps in the botanical record from such excavations.

**Figure 5.**
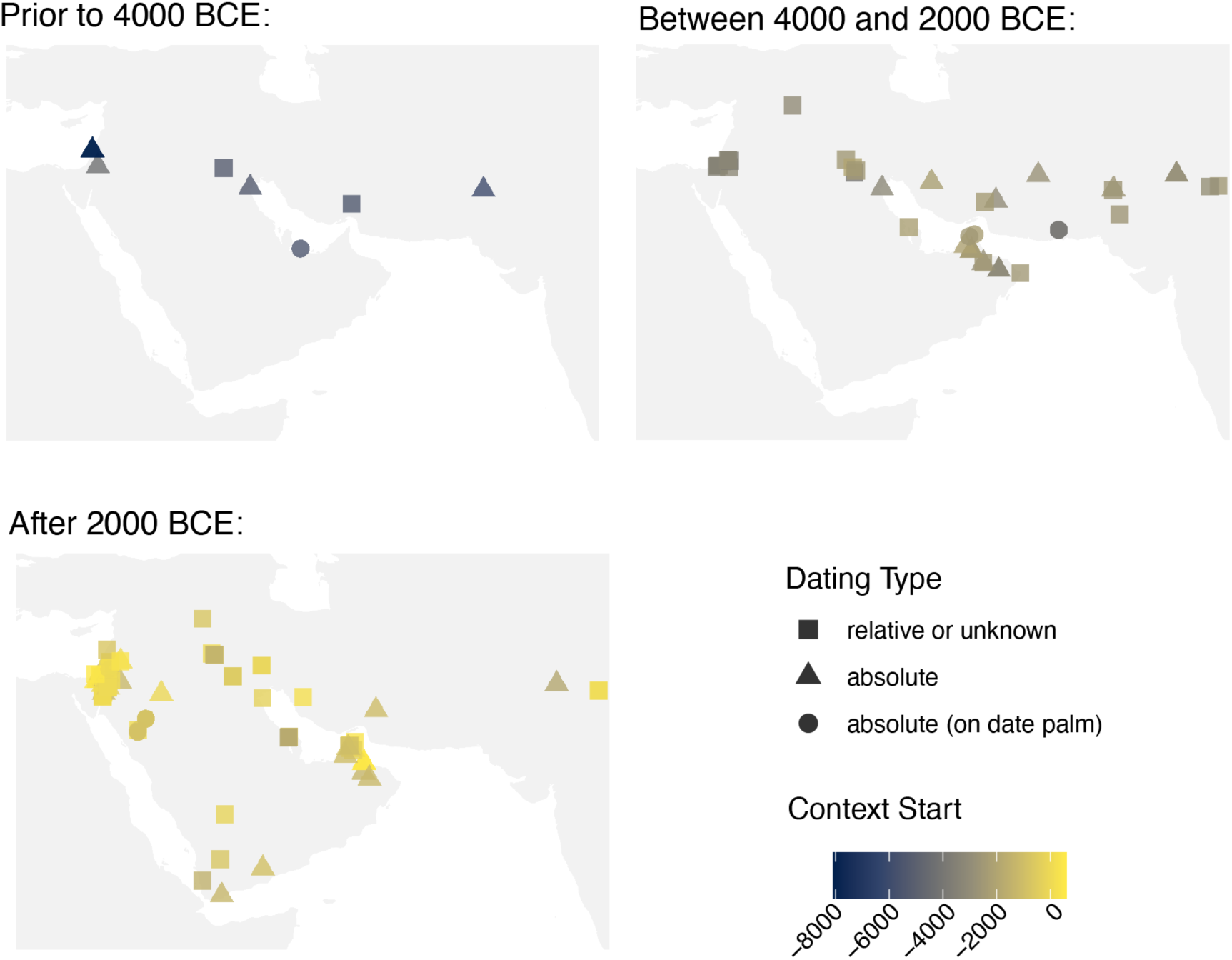
Geographical distribution of the 110 archaeological sites with *Phoenix* macroremains reported in the 123 reviewed sources. Colors correspond to the chronological starting point of each context; shapes denote the type and reliability of the dating.

## Discussion

In this study, we reviewed the literature and compiled archaeobotanical macroremains of *Phoenix* spp. in West and South Asia from prehistory to Late Antiquity. The findings were integrated into the web platform *DateBack*, which centralizes the data in structured tables and visualization tools designed to explore spatial and temporal patterns in the *Phoenix* archaeobotanical record. We used these records to propose a first interpretative synthesis on the early evidence of date use and cultivation across different regions of Asia—an initial step that can be further refined through more detailed future analyses and with the continued development of the platform.

### Synthesizing archaeobotanical evidence of *Phoenix*: Presenting *DateBack*, a platform for data compilation and visualization

Our literature review drew from 123 accessible sources. Despite our efforts toward comprehensive coverage, some references remained inaccessible due to language barriers, lack of digitization, or restricted availability. These challenges reflect broader issues in archaeobotanical research (Lodwick, 2019), and emphasize the importance of open science practices in ensuring data accessibility across disciplines (Wilkinson et al., 2016). By consolidating scattered records into an open-access platform, *DateBack* helps mitigate these challenges, providing a centralized and structured resource for future studies. Additionally, researchers can contribute new data or corrections via the provided contact email, ensuring the platform remains dynamic and continuously refined. All updates are tracked through a versioning system, which documents changes to both the data tables and the interface.

Our reliance on original sources may have led to the omission of some data, yet this approach ensured the accuracy and reliability of the recorded dataset by avoiding errors often introduced through secondary citations. For instance, we found that *Phoenix* remains were reported from Ain Ghazal (present-day Jordan), yet no such evidence appears in the original publication (Rollefson et al., 1985). By systematically verifying each original reference, we corrected such misattributions, ensuring that misreported data were not perpetuated in our dataset.

With this verified dataset, we integrated 154 entries from 110 sites across 15 countries into *DateBack*’s main dataset, alongside 74 radiocarbon-dated entries in a separate table. Compared to previous reviews, our dataset offers more standardized and detailed records. For instance, we systematically assigned geographic coordinates to each site, facilitating spatial analyses and visualizations, and recorded excavation years, which may serve as an indirect measure of data reliability, as identification methods and counting methods may differ between older and more recent excavations (Marston et al., 2014; Cappers and Neef, 2012). Although previous reviews have recorded dating quality (e.g., Flowers et al., 2019), this study is the first to not only distinguish between absolute and relative dating but also systematically compile all radiocarbon dates performed on *Phoenix* remains within our review’s scope, enhancing chronological reliability.

Future developments are expected to improve the usability, transparency, accessibility, and analytical depth of the platform, reinforcing *DateBack*’s role as a key resource for archaeobotanical research. For instance, the present dataset does not yet include an assessment of data confidence, as suggested in previous studies (e.g., Fuller & Weber, 2005, discussed in Flowers et al., 2019). Implementing such a grading system in future versions, with the potential for community contributions, could allow users to distinguish between well-documented remains and those with ambiguous identification or uncertain dating. Additional metadata fields are planned to track re-examinations of remains, such as seed morphometric analyses or ancient DNA studies, with links to relevant publications, a functionality that is already under development. Documenting storage locations will facilitate further research and validation, while integrating photographs of remains, when available, will support identification and comparative analyses.

### Reassessing the earliest archaeobotanical evidence of *Phoenix* before 4000 BCE

The earliest archaeobotanical evidence of date palm (*Phoenix* sp.) prior to 4000 BCE, as recorded in the database, consists of nine entries from seven archaeological sites across the Persian Gulf region, the Levant, and Pakistan (Figure 5). However, as detailed below, the nature, preservation state, and contextual or chronological reliability of these remains vary significantly. Their interpretation is further complicated by unknown past distributions of *Phoenix* species, including wild date palms (Barrow, 1998; Gros-Balthazard & Flowers, 2021) and difficulty to differentiate them morphologically (e.g., Gros-Balthazard et al., 2016).

The earliest reported *Phoenix* evidence comes from Ohalo II (21000-17000 BCE) in the Levant, where a single “wood” fragment was recovered from a surface layer of the excavated campsite (Liphschitz & Nadel, 1997). Given the scarcity of material, uncertain depositional context, and lack of direct radiocarbon dating, this evidence remains highly uncertain and cannot be confidently attributed to this early period.

Additional finds from the Levant also present challenges in terms of dating and interpretation. At ‘Atlit (8100-7400 BCE), a waterlogged seed and a “wood” fragment were recovered, but their precise archaeological context is unclear, as sources report their presence in either a well or hearths (Galili et al., 1993; Kislev et al., 2004). Furthermore, the taxonomic identification is inconsistent: while the “wood” was identified as *P. dactylifera* (Galili et al., 1993), the seeds were later attributed to *P. theophrasti* (Kislev et al., 2004). Another site, the Cave of Treasure, has yielded numerous date seeds, interpreted as domesticated based on their size (Fuller, 2018). Presumably from the Chalcolithic layer (4250–4050 BCE), their attribution remains uncertain due to the absence of direct radiocarbon dating and the site’s later Roman-period occupation, as noted by the authors (Zaitschek 1961; Bar-Adon 1980).

On the eastern side of the date palm’s range, early evidence from present-day Iran and Pakistan presents similar dating uncertainties. At Mehrgarh, two date seeds were recovered from 6^th^ millennium BCE contexts (Costantini 1984; 1985), but their silicified state prevents direct radiocarbon dating. Another site, Tepe Gaz Tavila (5400–4800 BCE), has also yielded one carbonized date seed (Costantini, 1985). For both sites, beyond dating issues, the limited quantity of seeds, lack of associated charcoal evidence, and the suggested presence of wild date palms in the region (Fuller & Stevens, 2019) make it unclear whether the seeds were locally produced or introduced, and whether they came from wild or cultivated palms.

In contrast to the uncertainties surrounding those early *Phoenix* remains from the Levant, present-day Iran, and Pakistan, archaeobotanical evidence from the Persian Gulf region appears more chronologically and contextually reliable, with several sites yielding remains from the late 6^th^ to early 5^th^ millennium BCE. The least-documented case is Tell al-Oueli in present-day Iraq, where stem or petiole fragments were recovered from Late Ubaid contexts (5200-4350 BCE; Neef, 1991). While the archaeological context is well-documented, the remains are unquantified, their state of preservation unknown, and the absence of radiocarbon dating prevents confirmation of their exact age, altogether making their interpretation difficult.

Stronger evidence comes from further south with the site of Aş Şabīyah (5300-4900 BCE) in present-day Kuwait, where four mineralized date seeds were found (Beech, 2003). Although their mineralized state precludes direct radiocarbon dating, indirect dating based on associated materials supports their chronological placement (Parker, 2010), and morphometric analyses suggest they correspond to dates gathered from wild populations (Gros-Balthazard et al., 2017), aligning with this ancient timeframe.

The more securely dated remains come from Dalma 11, on Dalma Island in the southern Gulf, where two charred date seeds and seed imprints were discovered (Beech & Shepherd, 2001). The seeds, directly radiocarbon-dated to 5290-4940 cal. BCE and 4810-4540 cal. BCE, are the oldest yet reported for the species. The presence of Ubaid pottery, a material culture tradition originating in southern Mesopotamia, suggests that these date fruits may have been imported from the northern Gulf region, although their exact origin remains uncertain (Tengberg, 2012). Regrettably, their poor conditions prevent morphometric analysis from determining whether they were wild or cultivated.

Overall, these findings illustrate both the insights and limitations that archaeobotanical evidence provides in reconstructing early date palm use. In the Persian Gulf region, date consumption is securely attested by 5000 BCE, whereas elsewhere, the evidence remains ambiguous. Among these finds, no clear evidence of cultivation has emerged, as the remains that could be analyzed align with wild populations, while others are too poorly preserved for assessment.

### The transition to date palm cultivation during the 4^th^–3^rd^ millennia BCE

During the 4^th^ and 3^rd^ millennia BCE, our review revealed that the number of archaeobotanical records of *Phoenix* increases significantly with 39 recorded entries from 34 sites spanning the Levant, Mesopotamia, the Gulf and the Indus valley region. While earlier evidence remains ambiguous, finds from this period suggest a growing reliance on date palms, with indications of early cultivation emerging, particularly in southern Mesopotamia and the Persian Gulf region.

Archaeobotanical remains from southern Mesopotamia and the northern Gulf contrast with earlier findings, as they appear in significantly larger quantities and more consistently include a diverse range of anatomical parts, such as fibers and leaf fragments, alongside seeds. At Eridu (4000-3000 BCE), abundant charred seeds were excavated and considered domesticated by the author (Gillett, 1981) and others (e.g., Fuller & Stevens, 2019), though the absence of radiocarbon dating limits chronological certainty. In contrast, at Al-Khidr (3200-1200 BCE), dozens of seeds and leaf imprints on bitumen were recovered, with radiocarbon dating of the occupation providing a more secure chronological framework (Hajnalová et al., 2009). This substantial and varied assemblage suggests that date palms were actively managed and cultivated rather than merely exploited from wild populations.

This limited number of finds, largely due to geopolitical constraints that have restricted archaeological investigations in the region, likely leaves additional evidence undocumented. However, other sources complement the archaeobotanical record. Iconographic depictions, such as the Warka vase from Late Uruk (4^th^ millennium BCE), illustrate date palm gardens, while textual sources from the late 4^th^/3^rd^ millennium BCE attest to the presence of date palm groves where date palms were cultivated alongside other crops (Landsberger, 1967; Tengberg, 2012; Miller, 2016; Michel-Dansac & Caubet, 2013; Paszke, 2019). Together, these records strengthen the hypothesis of early cultivation in southern Mesopotamia and the northern Gulf region.

Archaeobotanical evidence from southern modern Iran and the Indo-Iranian borderland is limited, resulting in a fragmented and uneven picture of date palm management during the 4^th^ and 3^rd^ millennia BCE. Yet, at Konār Şandal (3^rd^ millenium BCE), the discovery of both seeds and vegetative parts alongside cereals and grape remains provides compelling evidence for date palm cultivation (Tengberg, 2012; Mashkour et al., 2013). This interpretation is further supported by depictions of date palms chlorite vases from the same site (Perrot & Madjidzadeh, 2005). Conversely, in southwestern Pakistan, date seeds from Miri Kalat and Shahi Tump (Tengberg, 1998; 1999b) were directly radiocarbon dated to the early 3^rd^ millennium BCE and identified as wild (undomesticated) through geometric morphometric analyses, though their classification as *P. dactylifera* or *Phoenix sylvestris* (L.) Roxb., 1832 remains uncertain (Ivorra et al., 2024). The contrast with Konār Şandal underscores how, within the same region, human populations engaged with date palms in diverse ways, ranging from reliance on wild stands to active cultivation. Aside from these discoveries, available data remain scarce (Costantini, 1985; Miller, 1982), largely due to limited excavations and geopolitical constraints. Notably, no archaeobotanical evidence has been recovered from the arid lowland of southwestern and south-central Iran, where conditions would have been suitable for date palm cultivation, prompting Tengberg (2012) to suggest that date palms may have been grown there as early as in Mesopotamia.

In eastern Arabia, apart from the two previously mentioned 5^th^ millennium BCE seeds from Dalma islands, which may have been imported from Mesopotamia (Beech & Shepherd, 2001; Tengberg, 2012), no date palm remains are found before the end of the 4^th^ millennium. At Hili 8, one of the earliest finds in the region (3000-2700 BCE), abundant seeds and carbonised vegetative parts were recovered alongside cereals, with the occupation layer securely dated by radiocarbon (Cleuziou 1982; 1997; Tengberg, 1998). Through the 3^rd^ millennium BCE, charred date palm seeds and vegetative fragments alongside cereals appear at several sites, including Bāt (Tengberg, 1998; Deckers et al., 2019) and Qal’at al-Bahrain (Tengberg & Lombard, 2001). Although not all finds are radiocarbon-dated, their abundance and association with other cultivated species suggest that date palm cultivation was established in eastern Arabia by the 3^rd^ millennium BCE.

Further east, in the Indus Valley region and its eastward surroundings, within the context of the Harappan culture (second half of the 3^rd^ millennium BCE), date palm remains have been identified at several sites, among which are notably Mehrgarh, with numerous palm branches and stem fragments likely used for construction (Jarrige et al., 1995), and Nausharo, where abundant seeds were recovered (Costantini, 1990a). These findings, associated with the broader diffusion of crops such as cereals, pulses, and cotton (*Gossypium* sp.) into the region, have been interpreted as evidence of date palm cultivation by Tengberg (2012). In this context, it is worth noting that a few seeds were also recovered at the sites of Balu and Kunal, located in the eastern extent of the Indus cultural sphere (Saraswat & Pokharia 2002, 2003). Although their precise identification remains uncertain—whether they correspond *to P. dactylifera* or *P. sylvestris*—these remains are notable, especially given the scarcity of archaeobotanical evidence for date palms in this part of South Asia. As mentioned above for Miri Kalat and Shahi Tump in the Indo-Iranian borderland, these seeds may reflect exploitation of either wild date palms—possibly local populations—or another *Phoenix* species (*P. sylvestris*), constituting particularly relevant findings in light of ongoing research into early human-*Phoenix* interactions.

In the Levant, numerous sites have yielded date palm remains from the 4^th^ and 3^rd^ millennia BCE. However, as with the presumed earlier finds discussed above, these are limited to either seeds or charcoal and appear in small quantities. Only Qumrân, cave 24, has two secured direct datings providing the earliest confirmed evidence of date fruit consumption in the region (2920-2657 cal. BCE and 2897-2618 cal. BCE; Patrich, 1994; Liphschitz & Bonani, 2001; Sallon et al., 2020). Based on seed size, these were interpreted as wild (Liphschitz & Bonani, 2001), and other scholars suggest the presence of wild date palm in the region (Zohary & Spiegel-Roy, 1975), supported by *Phoenix* pollen evidence through the Holocene (e.g., Litt et al., 2012). However, taxonomic uncertainties persist, and re-examining charcoal assemblages analyzed decades ago using updated anatomical criteria (Bouchaud et al., 2012; Thomas, 2013) could help clarify identifications, particularly distinguishing between stipe and petiole fragments or refining genus-level classifications (e.g., Bronze Age charcoal fragments from Jericho; Western, 1971). While the recurrent finds reviewed here suggest the presence and use of date palms in the Levant during the Chalcolithic and Bronze Age, we consider the current evidence to be inconclusive regarding local cultivation, as proposed in previous studies (Langgut & Sasi, 2023; Fuller, 2018). Additional well-contextualized and larger botanical assemblages would help further evaluate this hypothesis. Our findings, in light of the literature, suggest that date palm cultivation emerged in southern Mesopotamia and the northern Gulf by the 4^th^ millennium BCE, while in eastern Arabia and the Indus Valley, the first evidence of phoeniciculture appears in the 3^rd^ millennium BCE. This pattern aligns with broader agricultural developments in the region. Cereal agriculture is known to have been introduced to eastern Arabia (Cleuziou, 2004), and trade networks linked Mesopotamia, the northern shore of the Persian Gulf, and the Indus Valley, with the Gulf serving as a hub for exchange and agricultural transmission (Beech & Shepherd, 2001; Tengberg, 2012). Given that the earliest secure evidence for date palm cultivation comes from southern Mesopotamia and the northern Gulf region, it is plausible that, like cereals, date palm cultivation in eastern Arabia—and perhaps also in the Indus Valley—was introduced rather than developing independently. In contrast, while consumption evidence exists in the Indo-Iranian borderland and the Levant, cultivation remains uncertain during this period.

### The late expansion of date palm cultivation throughout Southwest and South Asia in the 2^nd^ and 1^st^ millennia BCE

Although date palm cultivation may have begun earlier, the first secure evidence in the Levant comes from the Iron Age, with well-documented finds at two sites: Khirbet An Nahhas (1199-844 BCE, Baierle et al., 1989; Engel, 1993a) and Khirbat al-Jariya (1100-900 BCE; Liss et al., 2020; Stroth et al., 2023). Their dating relies on radiocarbon analysis, including on date palm remains for the later, and both seeds and charcoal were found in substantial quantities.

Unlike the Levant, where older remains exist but are often difficult to interpret due to uncertain dating and taxonomic challenges, in northwestern Arabia, the earliest securely dated remains also represent the oldest known evidence of date palm cultivation in the region. At Tall al-Sālimīyyah in the al-’Ulā region, seeds, stem, and petiole fragments have been radiocarbon-dated to 1215–1015 cal. BCE, supporting the hypothesis of local cultivation (Rohmer et al., 2022). At the same period, charcoal and seeds, some of them radiocarbon-dated, are attested at the neighboring site of Taymāʾ (Dinies et al., 2016; Tourtet et al., 2021).

In southern Arabia, the dataset compiled in *DateBack* is limited due to both geopolitical constraints limiting archaeological excavations and restricted access to original reports. While the potential existence of date seeds is reported from ar-Raqlah 1 (Costantini, 1990b), these occur only as pottery imprints, and thus were not included in *DateBack* as they fall outside the scope of our review. As a result, our dataset is constrained to four sites. At Şabir, remains described generically as “palm” have been reported from the 2^nd^ millennium BCE (Vogt & Sedov, 1998; Görsdorf & Vogt, 2001), but without clear taxonomic identification their attribution to *Phoenix dactylifera* remains uncertain, and direct radiocarbon dating is lacking. At Al-Kashawba’, a unique sounding revealed the presence of charred date palm seeds supposedly dated to the early 1^st^ millennium BCE, but no more detailed information is available (Phillips, 2007; de Moulins & Phillips, 2009). In contrast, securely identified desiccated date seeds from Raybūn (Levkovskaya & Filatenko, 1992) and charred seeds and imprints from Barāqish (de Maigret et al., 1986) provide more reliable evidence for date consumption by the 1^st^ millennium BCE. Since early to mid-1^st^ millennium BCE inscriptions explicitly mention palm groves (reviewed in Schiettecatte, 2013), it is highly likely that date palms were already under cultivation during this period.

In central Arabia, data are limited to the unquantified remains from Qaryat al-Fāw, a site occupied approximately 2000 years ago (Muṣṭafā Ḥasan et al., 2019). More material from the contemporaneous site of al-Yamāma (Schiettecatte et al., 2015) remains unpublished (Bouchaud, unpublished; Chambraud et al., ongoing). The currently available data thus suggest a late emergence of date palm gardens in central Arabia compared to neighboring regions, aligning with the relatively late development of urban oases in the area (Schiettecatte & al-Ghazzi, 2016). However, the lack of archaeobotanical studies makes it difficult to determine whether earlier cultivation may have occurred but remains undocumented.

In the continental part of northeast Arabia, archaeobotanical data are notably lacking, probably due to the limited number of large-scale archaeological projects. However, archaeobotanical analyses on the Classical city of Thaj (present-day Saudi Arabia), dated to the 7^th^ century BCE-7^th^ century CE, revealed the presence of date remains (seeds, fruits, stem and petiole) but their study and dating are still on going (Dabrowski, ongoing).

## Conclusion & Prospects

In this study, we have developed the *DateBack* platform, a centralized resource designed to facilitate research on the history, domestication, and diffusion of the date palm (*Phoenix dactylifera* L.). By rigorously compiling and standardizing archaeological and archaeobotanical data, the platform enables a refined assessment of the spatial and temporal patterns of date palm use and management.

Our findings confirm that the earliest secure evidence for date consumption is concentrated in the Persian Gulf, while evidence from other regions, particularly the Levant, remains ambiguous due to uncertain contexts, taxonomic issues, and a lack of direct dating. Date palm cultivation appears established in southern Mesopotamia and the northern Gulf by the 4^th^–3^rd^ millennium BCE and may have developed as early as in southern Iran, though the scarcity of archaeobotanical studies prevents confirmation. In the Indus Valley region and eastern Arabia, cultivation is attested by the 3^rd^ millennium BCE, and connections with the Gulf may indicate a diffusion process rather than independent emergence.

In the Levant and northwestern Arabia, secure published evidence for cultivation appears at the end of the 2^nd^ millennium BCE, while in southern Arabia, it is not attested until the 1^st^ millennium BCE. The late emergence of date palm cultivation in various regions of the Arabian Peninsula seems to result from a combination of regional factors. In northwestern Arabia, available data suggest a distinct regional agricultural pattern where the cultivation of date palm is integrated to pre-existing agricultural systems (Dinies et al., 2016; Rohmer et al., 2022; Chambraud, unpublished). Elsewhere, particularly in southern and central Arabia, limitations stem primarily from the scarcity of archaeobotanical studies or challenges in interpreting existing evidence. Additionally, the lack of Bronze Age contexts narrows the chronological perspective, and restricted access to reports and publications continues to hinder data collection.

While this study has reviewed date palm remains from earliest evidence through Late Antiquity, their interpretation has primarily focused on early date consumption and emergence of cultivation in South and Southwest Asia. Further research is needed to explore additional questions, including continuity—whether date palm cultivation was consistently maintained in certain regions or periodically abandoned—and diffusion routes. In addition, morphometric analyses of well-preserved seeds could help assess their wild or domesticated status and refine the understanding of date palm consumption and cultivation in different regions. Expanding archaeological research, particularly in data-poor areas, would also be essential to filling critical gaps in current knowledge. The *DateBack* platform is designed as a dynamic and scalable resource, with future developments aiming to extend its scope to later periods (e.g., Islamic and beyond) and additional regions such as North Africa. Beyond macroremains, forthcoming updates may incorporate data on microremains (e.g., pollen, phytoliths), with initial work on this expansion already underway but not yet included in the current public version. Further iterations could also integrate impressions, textual and iconographic evidence, enhancing the interpretative framework. By fostering interdisciplinary collaboration among archaeobotanists, historians, and geneticists, *DateBack* will continue to refine our understanding of the cultural and agricultural significance of the date palm, adapting as new discoveries and methodologies emerge.

## Appendices

## Appendix 1 Structure and details of the Main Data Table fields

Appendix 1 - Structure and details of the Main Data Table fields.
| Field | Format and details |
| --- | --- |
| **Archaeological site** |  |
| Site | Matched with names from https://www.geonames.org/ when possible. If no match, used the name as in the original article. Multiple names may be provided for sites with different names (e.g., Al Baḥrīyāt (Isin)). |
| Type of site | From a limited list: port; settlement; town/city; burial site; villa; fort/castle (qasr); shipwreck; necropolis/cemetery; village; mine; public building; cave; campsite; monastery; craft area. |
| Excavation years | If available, the years during which the excavation took place. |
| Country | Present-day country of the archaeological site. |
| Latitude/Longitude | Decimal format. Gathered from https://www.geonames.org/ whenever possible or from Google Maps. |
| Region | Wide regions, for coloring plots. |
| **Chronological context** |  |
| Context(s) | Identification of the context(s) in which date palm remains were recovered (e.g., stratigraphic unit, locus, or area). |
| Start date/Finish date | The minimum and maximum dates assigned to the archaeological context. |
| Cultural period | Broadest period possible, to avoid ambiguity. |
| Dating quality | "Absolute" if the dating of the stratigraphic unit or the date palm remains themselves is based on radiocarbon (C14) dating; "Absolute on date palm" if the radiocarbon dating specifically pertains to the date palm remains; "Relative" otherwise, when dating is based on stratigraphy or other non-absolute methods. |
| **Remains** |  |
| Taxon (original) | As reported in the original study or paper. |
| Taxon (corrected) | Correction of the taxon to resolve ambiguities or address potential errors identified in the original study. "NA" if no correction applied. |
| Comments | Explanation of the taxon correction, including the reason and details of the change. |
| Anatomical part(s) | From a limited list: endocarp; flower; fibre; fruit/pericarp; leaflet; pedicel; perianth; petiole; seed; stem; stem/petiole (when the exact part is uncertain).<br>Possibility of multiple types of remain for a single entry. |
| Preservation | From a limited list: desiccated; roasted; charred; mineralized; waterlogged; silicified.<br>Possibility of multiple states of preservation for a single entry. |
| Quantity of remain | Free text providing the quantity or number of fragments recovered, where available. |
| C14 on date palm | Was radiocarbon-dating performed on a date palm remain ? |
| Reference(s) | Citation of the published source(s) that reported the remains, corrected their identification, or provided dating information, including links to the resource when available. |
| **Versioning** |  |
| Inclusion | Indicates the version number where the entry was added |
| Modification | Indicates the version number where the entry was modified (NA indicates no modification) |

## Appendix 2 Main Data Table summarizing the results of the literature review. Details and formats of the fields are provided in Appendix 1. For entries with radiocarbon dating performed on date palm remains, corresponding results are available in Appendix 3

## Appendix 3 Radiocarbon data table for date palm remains. This table documents radiocarbon analyses of date palm remains, linking each analysis to its site and context. It includes specific fields such as laboratory codes, anatomical parts, preservation states, raw ^14^C ages (with uncertainty), and calibrated age ranges

## Acknowledgements

We would like to acknowledge Naomi Miller, Margareta Tengberg, Guy Bar-Oz, Daniel Fuks and Laurent Bouby for their valuable contributions to the archaeobotanical review. Special thanks to Yann Bourgeois and Vincent Battesti for their insightful discussions and comments on the manuscript. We also appreciate Anil K. Pokharia, Stéphanie Thiébault, Michael Purugganan, and Jonathan Flowers for their assistance in accessing some key documents. Additionally, we would like to thank the IRD documentation team in Montpellier, particularly Aure Franceschi and Hanka Hensen, for their help in locating references, as well as the librarians at NYU for their support.

We are grateful to Claudia Speciale, Claudia Moricca (Reviewer 1), and the anonymous Reviewer 2 for their constructive and insightful comments, which have helped us improve the clarity and scope of the manuscript, and for helping us to access some documents that we had not been able to find.

The *DateBack* logo has been designed by Emeline Pobelle (https://www.malt.fr/profile/emelinepobelle).

AI tools, specifically ChatGPT and DeepL, were used to assist in polishing and refining the text throughout the manuscript, and in writing the code of the web application.

A preprint version of this article has been peer-reviewed and recommended by PCI Archaeo (https://doi.org/10.24072/pci.archaeo.100603).

## Data, scripts, code, and supplementary information availability

The structure of the data tables is licensed under CC-BY, and the contents of the data tables are licensed according to the original datasource or CC-BY.

Data are available online. The general repository can be found at https://dataverse.ird.fr/dataverse/dateback. It includes the main table (Appendix 2: https://doi.org/10.23708/4ZHYZU) and the table with the radiocarbon dating (Appendix 3: https://doi.org/10.23708/C5N9CM).

Scripts and code are available online in a git repository: https://forge.ird.fr/diade/besseiche/dateback (https://doi.org/10.23708/KNQIN1 for v.1.0). All codes are released under the license GNU GPL v3.

The web application with the currently described version of the data table is hosted by the French “Institut Français de Bioinformatique” (IFB) at https://cloudapps.france-bioinformatique.fr/dateback

## Contribution of each author

The design of the data table was primarily carried out by MB, EC, VD, CB, MGB, with additional contributions from EB. The literature review and data entry were mainly performed by EB, EC, VD, CB, MGB, with contributions from MB. The curation of the data table was led by MB and MGB. The design and development of the web application was primarily handled by MB, with input from FS and MGB. The manuscript was primarily written by MB and MGB, with revisions and input from all other co-authors.

